# Variation of sexual dimorphism of wing shape in the family Dolichopodidae (Diptera)

**DOI:** 10.1101/2022.02.25.481985

**Authors:** M.A. Chursina, O.O. Maslova

## Abstract

Although sexual dimorphism manifestations are widespread in the family Dolichopodidae, a detailed characterization of their phylogenetic significance is lacking. Purpose: in order to study the distribution patterns of wing sexual dimorphism, we analysed 57 species from 17 genera of 9 subfamilies. Methods: A comparative analysis of the evidence, obtained by geometric morphometry and molecular data, allowed us to assess the phylogenetic signal in the sexual dimorphism of the wing. Results: The results of the study confirm the presence of diverse patterns of sexual variability in the wings of this family. More often, females have larger wings with blunted apexes, whereas males are characterized by a more pointed apex. In some cases, the larger size of females’ wings is associated with an increase in body size, while in other cases, differences in shape and size can be explained by differences in behavioural and life patterns. Although there exists a general pattern of sexual dimorphism, its features differ even in closely related species. Conclusion: the absence of a significant phylogenetic signal in seven out of nine studied wing points indicates that the sexual dimorphism in form evolved, at least partially, in each of the studied species.

---

Sexual dimorphism is a phenomenon frequently encountered in the Dolichopodidae family. The most frequent characters of sexual dimorphism are various modifications of tarsi (dense pubescence, distention and protrusion, colour changes), wings (colour changes of the wing membrane, thickened costa), postpedicel elongation, and modifications of arista (swelling or protrusion). Such indicators are usually used for diagnostics.

However, more impalpable distinctions between females and males, such as wing shape, are characteristic of the family, and the types of sexual dimorphism of wing shape change from species to species [1, p. 515]. For example, it is found that *Argyra* Macquart, 1834 males perform a mating dance in front of females [2, p. 11], and *Poecilobothrus nobilitatus* (Linnaeus, 1767) males exhibit aggressive demonstrations and chases in rivalry for females [3, p. 602]. Although behavioural traits are considered more evolutionarily labile, they also often carry a significant phylogenetic signal [4, p. 740; 5, p. 7]. In some cases, the wing shape variability can be caused by considerable differences in the body size of females and males, such as in the *Rhaphium appendiculatum* Zetterstedt, 1849.

The wide variety of sexual dimorphism of the wing shape suggests intensive selection. Along with traditional morphological and molecular traits, signs of sexual dimorphism are also a resource for phylogenetic constructions, although such studies are much rarer. Thus, a phylogenetic signal in the sexual dimorphism of the wing shape is evident in the *Drosophila* Fallén, 1823 species [6, p. 110]. And what is interesting is the phylogenetic reconstruction of the development of elongated ocelli in males in the family Diopsidae [7, p. 1373].

On the other hand, similar manifestations of sexual dimorphism often occur in non-closely related species. Examples include the formation of an elongated exoskeleton in cheese flies and nereid [8, p. 602], wing spots in fruit flies [9, p. 322], and protrusion and distention on the legs and other body parts in the Diptera of various families [10, p. 143]. Therefore, we can expect that some genetic factors play an essential role in forming a specific pattern of sexual dimorphism, which results in a more pronounced convergence in the morphological characters of nonrelated species than can be explained from a functional point of view.

The analysis of molecular data, together with the sexual dimorphism characters of wing shape, will allow us to consider evolutionary trends of sexual dimorphism, reconstruct ancestral forms, and possibly clarify some controversial points of the phylogenetic tree Dolichopodidae. In the current study, we analysed the phylogenetic signal of sexual dimorphism in the wing shape to reveal patterns of distribution between subfamilies and genera.

## Materials and methods

In total, 5874 specimens of wing of 57 species of 17 genera belonging to 9 subfamilies were studied (Table 1). we used individuals that we collected during 2013–2021 as well as those from the collection of the Department of Ecology and Systematics of Invertebrates, Voronezh State University (Voronezh, Russia).

**Table 1.**
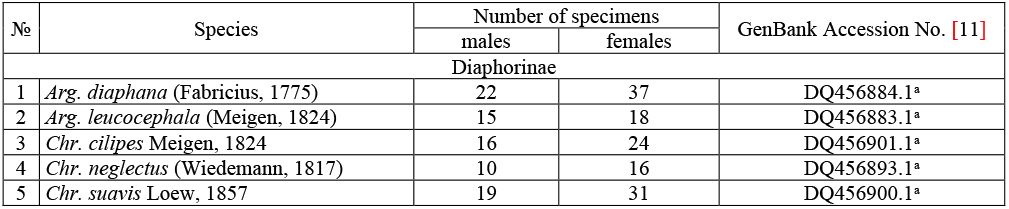
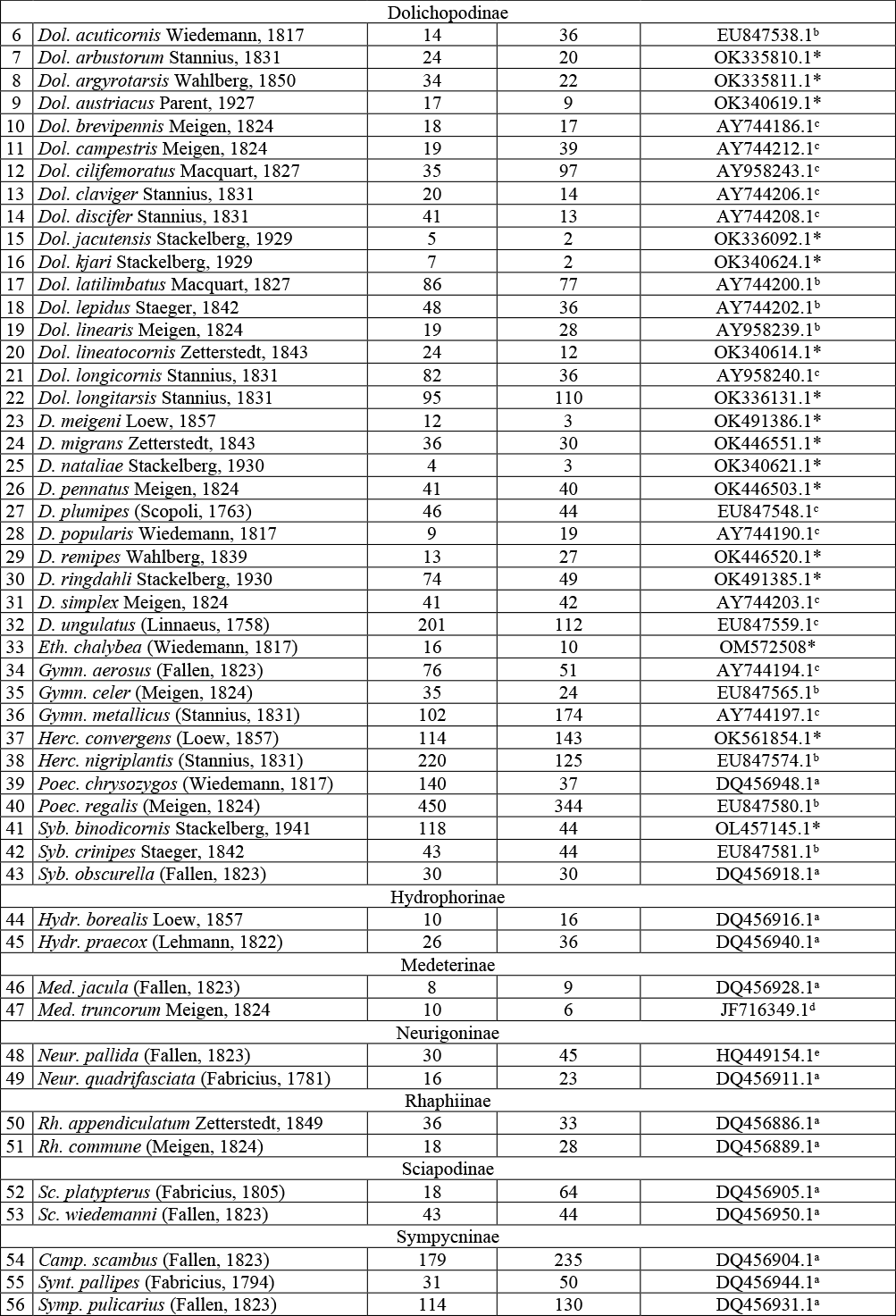
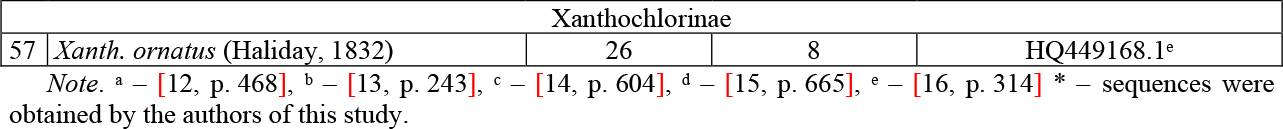
Studied species

The analyzed molecular matrix included molecular sequences of the mitochondrial gene encoding cytochrome c oxidase (COI) (810 characters). The study included both sequences previously deposited in GenBank (GenBank, 2021) and sequences carried out especially for this study by the Sintol Enterprise (Russia). In total molecular sequences of 57 species was studies. Amplification and sequencing were performed using the methods and primers described in previous studies [12, p. 455; 14, p. 605]. The sequences were aligned manually using BioEdit multiple alignment software [17]. Phylogenetic reconstruction was carried out using the minimum evolution method (ME) in MEGA software [18]. Reliability of inner branches was estimated by the bootstrap method based on 1000 pseudoreplicates.

Wings were digitized at 9 landmarks (fig. 1). Each landmark has been digitized using TpsDig-2.32 software [19].

**Figure 1.**
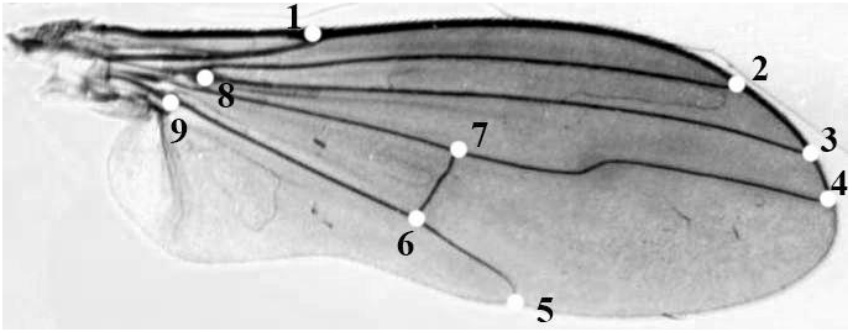
Wing and landmarks positions

For comparing overall wing size among different populations, we used the isometric estimator known as centroid size, which is defined as the square root of the sum of the squared distances between the center of the configuration of landmarks and each separate landmark [20, p. 56]. Shape variables were obtained through the Generalized Procrustes Analysis [20, p. 106]. Then, analysis was carried out using the methods of multivariate statistical analysis in software MorpholJ [21].

Canonical variate analysis was used to determine the most important differences between sexes, and the obtained canonical coefficients for each landmark were used in further analysis. To construct a dendrogram demonstrating the similarity of patterns of wing shape sexual dimorphism, the method of unweighted pair group method with arithmetic mean was used. The reliability of internal branching was assessed using bootstrap analysis with 1000 replicas. The statistical significance of pairwise differences in mean shapes of males and females was analysed using permutation tests (10 000 rounds) with Procrustes distances (PD) [21]. The allometric component of the wing shape variability is estimated using regression analysis of shape variables for centroid size.

The phylogenetic signal of wing sexual dimorphism was assessed in two ways. First, the phylogenetic tree (Figure 2) was superimposed on the space of shape variation, and then the hypothesis that the phylogenetic signal was absent was tested using a permutation test with 10 000 integrations. The main components of the shape variability were substituted into the nodes of the phylogenetic tree. The p-value was calculated as the fraction of permutations that lead to the length of the tree, which is equal to or less than that observed for the original data [6, p. 9].

**Figure 1.**
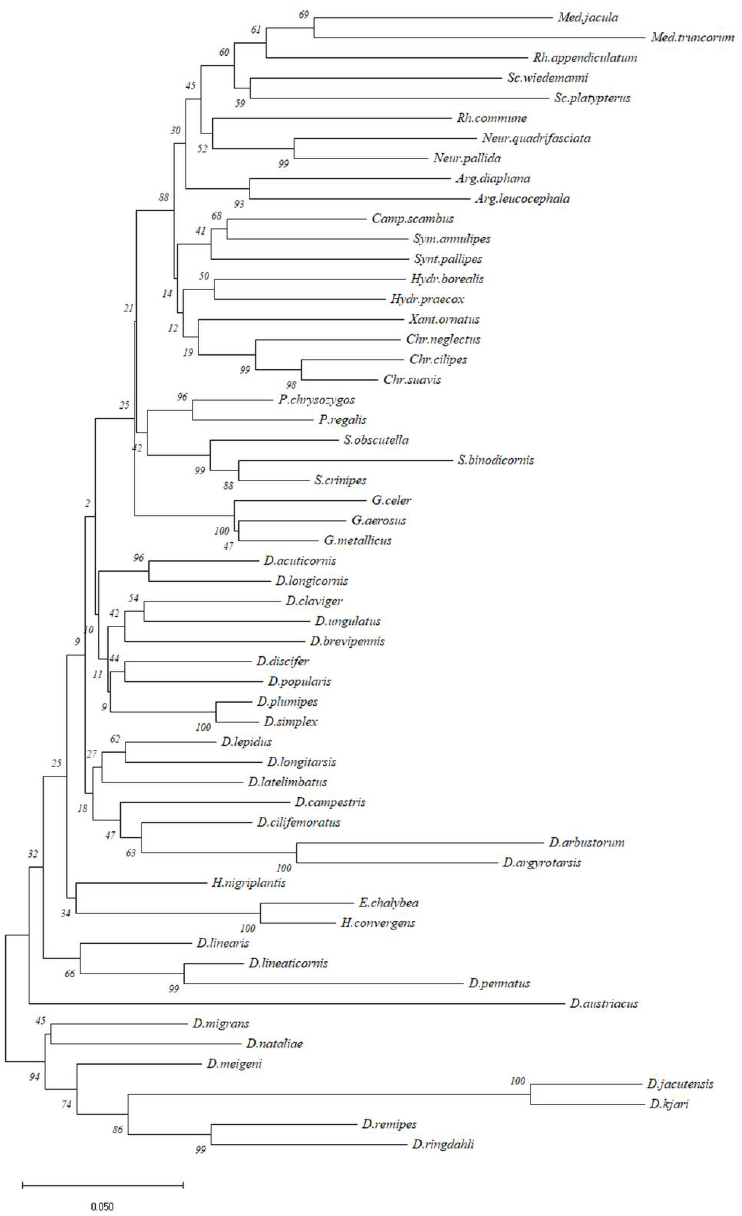
ME tree, obtained from COI sequences. Values of bootstrap support from 1000 pseudoreplicates are depicted above nodes

Secondly, as a measure of phylogenetic signal of legs morphometric characters, we used Pagel’s lambda (λ) [22] and Blomberg K-statistic [23]. To calculate Pagel’s lambda, the *phylosyg* function *phytools* package [24] was used in R environment [24]. Blomberg K-statistic also takes values from zero to one, but if the phylogenetic signal is very high, then K-statistic can rise over one. To calculate Blomberg K-statistic, the *Kkalk* function *picante* package was used in R environment [25]. For testing purpose the indications of differences of the metric from 0, a p-value was obtained by randomizing the trait data 1000 times.

## Results

The ANOVA demonstrated that the following factors have a significant effect on sexual dimorphism of wing size: “subfamilies × sex” (F = 3,5; df = 8; P = 0,0005); “genera × sex” (F = 17,5; df = 16; P < 0,0001) and “species × sex” (F = 12,0; df = 56; P < 0,0001). This means that significant differences are observed in the sexual dimorphism of wing size between subfamilies, between genera and between species. Moreover, in 44 cases out of 57, the females wing size exceeded the males wing size. In the species *Argyra, Chrysotus, Neurigona, Rhaphium, Sympycnus, Syntormon*, and *Xanthochlorus*, female wings were larger than those of males. In other species, both situations were encountered.

The smallest sexual difference in size was observed in the species *Campsicnemus scambus*, the largest in the species *Hydr. borealis, Rh. commune*, and *Dol. argyrotarsus* (female wings are larger than male wings), as well as *Syb. crinipes* (male wings are larger than female wings). Among the subfamilies, the largest variation of the difference in wing sizes was characteristic of the Dolichopodinae, the smallest mean value was observed in the subfamilies Medeterinae and Sciapodinae, and the largest in Rhaphiinae.

Differences in sexual dimorphism of wing shape were also significant between subfamilies (Wilks’ Lambda = 0,82; F = 20,3; df = 112, 81806,51; P < 0,0001), between genera (Wilks’ Lambda = 0,39; F = 51,5; df = 224, 123435,8; P < 0,0001) and between species (Wilks’ Lambda = 0,07; F = 42,9; df = 784, 157298,4; P < 0,0001).

The most pronounced sexual dimorphism in the form of a wing was observed in the species *Xanth. ornatus* (PD = 0.107; P < 0.0001) and *Arg. diaphana* (PD = 0.117; P < 0.0001), the least pronounced in *Gymn. aerosus* (PD = 0.006; P < 0.02) and *Herc. convergens* (PD = 0.006; P < 0.001). Of the subfamilies, the largest variation in PD values was characteristic of Sympycninae, the smallest average PD value was observed in the subfamilies Medeterinae, Hydrophorinae, and Rhaphiinae, and the largest in Sympycninae and Sciapodinae.

The differences in the sexual dimorphism of wing shape most often consisted in the displacement of Landmarks 3 and 4 along the x-axis, as well as Landmarks 5 along the y-axis, which, in the general case, led to the formation of a more elongated wing with a sharp apex in males and a more rounded wing with a blunt apex – in females.

According to the UPGMA-dendrogram, built on the basis of the canonical coefficients of sexual dimorphism, the most similarity in the sexual dimorphism of the wing shape was shown not always by phylogenetically related species. A similar shape dimorphism has been shown for the following species: *Dol. longitarsis* and *Dol. ungulatus* (bootstrap index BS = 78), *Dol. austriacus* and *Dol. lineaticornis* (BS = 50), *Dol. acuticornis* and *Gymn. aerosus* (BS = 53), *Syb. binodicornis* and *Sc. platypterus* (BS = 67), *Arg. leucocephala* and *Xanth. ornatus* (BS = 54). The sexual dimorphism of the wing shape of the *Medetera* species was clearly different from the other species of the family (Figure 3).

**Figure 3.**
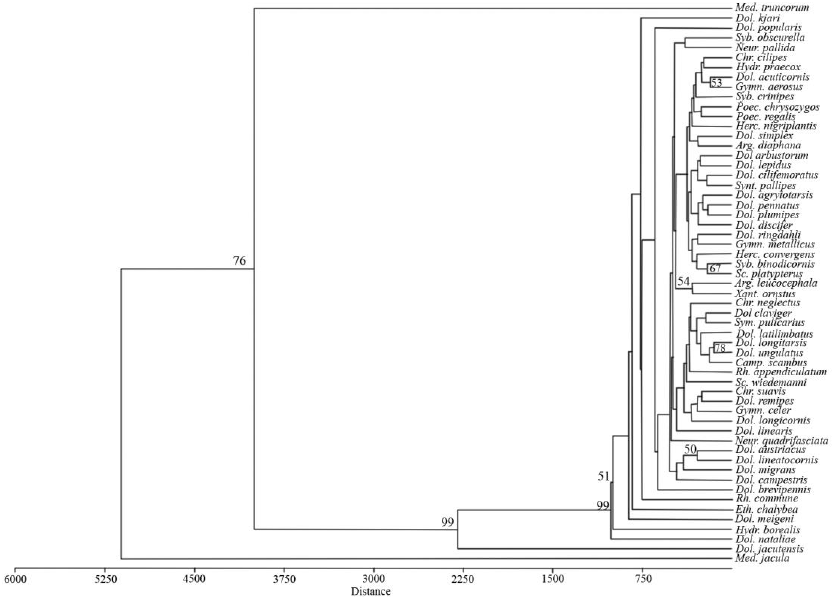
Results of UPGMA cluster analysis of the canonical coefficients of sexual dimorphism of dolichopodid

It should be noted that the allometric component of sexual dimorphism of the wing shape in most species is expressed insignificantly. The greatest percentage of shape variability associated with the sexual wings size difference was found in the following species: *Syb. obscurella* (40.9%; P < 0.0001), *Dol. meigeni* (37.5%; P < 0.0001), *Dol. austriacus* (34.6%; P < 0.0001), *Dol. kjari* (33.6%; P < 0.0001), *Arg. diaphana* (25.7%; P < 0.0001), *Neur. quadrifasciata* (25.7%; P < 0.0001). The species that showed the largest differences in wing size between females and males did not show a high percentage of allometric variation in shape.

The length of the consensus tree combining the initial molecular data and data on the wing shape changes was 0.1082 (in units of squared Procrustes distance) (Figure 4). The permutation test produced an equal or a longer tree in most cases (P < 0.0001), thus confirming the presence of a phylogenetic signal in interspecific variation of sexual dimorphism in the wing shape.

**Figure 4.**
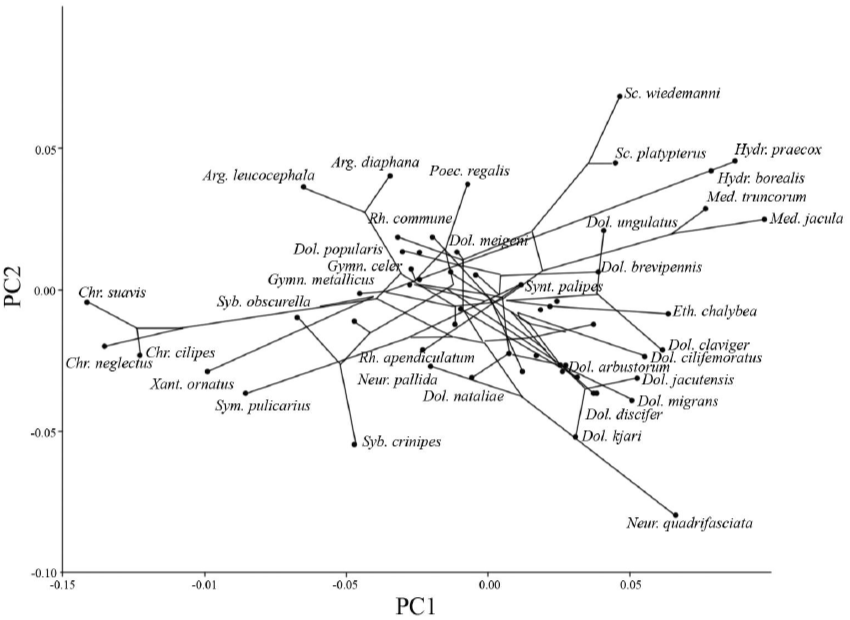
Schema fragment of the changes in the wing shape related to sexual dimorphism mapped onto phylogeny: the first (29.2%) and second (17.8%) principal components of variation

In the Procrustean distance between the wing shapes of females and males, the minimum phylogenetic signal was observed: λ = 0.00007, P = 1; K = 0.69, P = 0.15. The most significant phylogenetic signal was found for the canonical coefficients of landmarks X1 (λ = 0.99, P = 0.05; K = 1.19, P = 0.003), Y1 (λ = 0.99, P = 0.007; K = 1.31, P = 0.004), Y2 (λ = 0.99, P = 0.02; K = 1.23, P = 0.005).

## Discussion

Most of the studied species of Dolichopodidae showed a significant sexual dimorphism of wing shape and/or size, with an insignificant influence of allometry on shape variability. However, the sexual dimorphism of wings in the family is heterogeneous: species of one subfamily showed that the wing size of females exceeds that of males and vice versa; besides, we can distinguish species with significant sexual differences in both wing shape and size (*Arg. diaphana*), species with insignificant sexual differences both in shape and size (*Camp. scambus*), and species with significant differences in wing size and insignificant differences in shape (*Rh. commune*), and also species with slight sexual differences in wing size, but high differences in shape (*Arg. leucocephala, Eth. chalybea*). This means that different species are influenced by various selection factors, which may act together or independently on both sexes.

Studies show that female insects are more often larger than males because of the high correlation between body size and fecundity [26]. This may explain that in most cases, the wings of female dolichopodids are larger than those of males since wing size directly correlates with body size. This regularity is well documented in *Rhaphium* species, where differences in body sizes of females and males are maximal. At the same time, sexual dimorphism of wings is shown in differences of size, but not of form (*Rh. commune*), or available differences in form are partially explained by allometry (*Rh. appendiculatum*).

In other cases, when males had larger wing sizes, this could be explained by other factors, e.g., more significant load on males’ wings due to different behavioural patterns: fights between males, and peculiarities of mating dance (for example, in *Poec. regalis* males).

Differences in the wing shape of males and females differed in each case but more often consisted in the displacement of 3, 4, and 5 landmarks, i.e., the change in the distal wing part, while the base remains unchanged. The proximal region of the wing is most susceptible to changes, both in the case of sexual, interspecies, and intraspecies variability [1, p. 695].

Regardless of the factors influencing sexual dimorphism of wings, our results show that patterns of sexual dimorphism can differ even in closely related species, since even species from different subfamilies turned out to be close in form. This is probably because, in each species, sexual dimorphism of wing shape and size results from complex interactions between several factors of selection that depend on the specific biology, genetic and ecological features, and ontogenetic history of each sex.

Although sexual dimorphism of wing shape appears to be somewhat dependent on common ancestry (the overall phylogenetic signal of sexual dimorphism was reliable), the absence of a significant phylogenetic signal for seven out of nine studied wing points indicates that sexual dimorphism of shape evolved, at least in part, in each studied species.

## Conclusion

Our study demonstrates that interspecific differences in the sexual distinction of wing shape in dolichopodids are most often nonallometric and do not depend on phylogenetic relationships between species. These differences are likely the result of a complex interaction of intra-sex competition and other types of selection acting with different intensity in each sex and on several interrelated characteristics, such as body size, wing size, and shape. Overall, the present study results demonstrate that the mechanisms responsible for the emergence of sex differences in wings can form different and complex patterns of sexual dimorphism in the family.

***The work was funded by RFBR and NSFC according to the research project № 20-54-53005***.

